# Exploring Survival Models Associated with MCI to AD Conversion: A Machine Learning Approach

**DOI:** 10.1101/836510

**Authors:** Jorge Orozco-Sanchez, Victor Trevino, Emmanuel Martinez-Ledesma, Joshua Farber, Jose Tamez-Peña

## Abstract

Several studies have documented that structural MRI findings are associated with the presence of early-stage Alzheimer Disease (AD). However, the association of each MRI feature with the rate of conversion from mild cognitive impairment (MCI) to AD in a multivariate setting has not been studied fully. The objective of this work is the comprehensive exploration of four different machine learning (ML) strategies to build MRI-based multivariate Cox regression models. These models evaluated the association of MRI features with the time of MCI to clinical AD conversion. We used 442 MCI subjects from the Alzheimer’s disease Neuroimaging Initiative (ADNI) study. Each subject was described by 346 MRI features and time to AD conversion. Cox regression models then estimated the rate of conversion. Models were built using four ML methodologies in a cross-validation (CV) setting. All the ML methods returned successful Cox models with different CV performances. The best model exhibited a concordance index of 0.84 (95% CI: 0.82-0.86). The final analysis described the hazard ratios (HR) of the top ten MRI features associated with MCI to AD conversion. Our results suggest ML exploration is a viable strategy for building and analyzing survival models that predict subjects at risk of AD conversion.

## I. INTRODUCTION

Dementia is one of the most common syndromes among the elderly [1]. According to Alzheimer’s Disease International, there are currently more than 50 million cases of dementia with an incidence of 10 million new cases each year. Alzheimer disease (AD) represents 60%-70% of the cases [2]. The most worrisome aspect of AD is the lack of effective therapy to control the disease; hence, between 2000 and 2015 the number of deaths caused by the disease increased by 123%. Developing an effective therapy is hampered by the complexity of aging, and the lack of a clear understanding of the etiology and pathogenesis of AD [3], [4]. Having a clear understanding of the AD process and stages is essential to develop effective therapies, while understanding the AD process requires an accurate diagnosis of the affected person.

Diagnosing of AD at early stages is complex. The most accurate AD diagnosis test requires the histopathologic evaluation of brain tissue via autopsy or biopsy [5]. In the absence of a biopsy, a typical person may be diagnosed with possible or probable AD, based on patient reports, cognitive observation, and symptomatology [6]. There are some useful clinical information such as Apolipoprotein E (APOE), that is a protein involved in the metabolism of fats in the body with a polymorphic structure that has three major alleles [7]. The fourth allele (APOE4) had been validated several times as a biomarker indicative of the risk of suffer Alzheirmer’s disease [8]. On the other hand, AD progression is slow. In early AD stages, patients do not have enough symptoms to be diagnosed with probable AD, but fall between the cognitive changes of aging and early dementia. This condition is known as Mild Cognitive Impairment (MCI) [9]. Therefore, an MCI diagnosis, in AD patients, is an intermediate stage between normal aging and clinical dementia, and only 33.6% of MCI subjects convert to clinical AD [10], [11], [12].

To address the lack of a definitive AD test, researchers have proposed the use of advanced image modalities – like magnetic resonance imaging (MRI) and positron emitting tomography (PET) – to support clinical diagnoses. Image-based findings associated with a disease process are called imaging-biomarkers [13], [14]. AD-related imaging-biomarkers have been found in MRI and PET, and have a clear association with the evolution and presence of AD [15], [16]. Furthermore, AD-related imaging-biomarkers have been associated with the conversion from MCI to AD [6], [17]–[20]. Therefore, MRI and PET are commonly used to monitor the progression of the disease and to detect the current stage of neuronal degeneration [21]. Although PET has the potential to directly visualize the AD-related molecular degeneration[22]–[24], this modality requires the use of radiopharmaceuticals, as well as facilities not as common as MRI [25]. In addition, MRI is safer than PET and quantitative MRI (qMRI) enhances MRI potential to detect minute anatomical changes related to the early AD process [26]. Hence, qMRI may become the standard screening procedure for early AD diagnosis. Therefore, studying the behavior of qMRI biomarkers associated with neurological degeneration is an important step in diagnosing and ultimately treating AD.

Discovering, characterizing and validating imaging-biomarkers associated with AD requires a well-designed study. The Alzheimer’s Disease Neuroimaging Initiative (ADNI) is a large study aimed to discover and test novel imaging-biomarkers [27]. ADNI has generated hundreds of research papers in this area, but most of them have used supervised classifications or statistical approaches for the characterization of MCI patients that presented with AD conversion. These research papers have been useful in discovering early imaging findings, but most of them have not evaluated effectively the time to AD conversion in their discovery efforts [28]–[30]. This evaluation is important because imaging biomarkers may be associated with an increase in time of conversion (low risk markers) or with a decrease in conversion time (high risk marker). Moreover, the magnitude of the time conversion increase or decrease (hazard ratio) may be specific for each marker. In this context, the multivariate Cox regression incorporates the time to an event, and evaluates the hazard-ratios (HR) of each potential-biomarker involved in the MCI to the AD conversion process. Therefore, Cox-based modeling has the potential to improve the understanding of imaging biomarkers associated with the AD process.

The limitation of Cox models in multivariate biomarker discovery is that model fitting requires selection of significant features, which commonly is done by either regularization or subset-selection; hence, most of the reported studies that have used Cox modeling have been limited to a small set of imaging biomarkers [31]–[33]. Statistical Learning (SL) and Machine learning (ML) approaches provide efficient and highly competitive solutions to the issues of regularization and subset selection. Embedded statistical learning like L1 regularization via de LASSO, allows the exploration of multivariate models composed on hundreds of features [34]. In addition, this technique allows subset-selection with the exploration of realizable Cox models from hundreds of features [35]. Model selection via the Bootstrap Step-Wise Model selection (BSWiMS), and Best Subset Selection (BeSS) are among two of the machine learning options readily available to researchers [35], [36]. Besides these approaches, feature selection (FS) is a common method used to build Cox models [37], [38]. The wide variety of methods available to researchers can make biomarker discovery a complex effort, especially when there is no clear choice of methodology for building/exploring survival models.

To overcome this limitation, we propose a unified approach for the study of Cox models in an ML setting. The approach is based on repeated cross-validating ML/SL methods using exactly the same training-testing sets across methods. The ML implementation evaluates LASSO, BSWiMS, BeSS, and Univariate Filtering for building suitable survival models. Thus, at the end of the repeated CV, a fair method comparison and a comprehensive evaluation of the role of each potential biomarker inside a Cox survival model is provided.

The main goal of this paper is the application of a unified approach to cross-validate Cox regression models for the exploration of survival-based analyses of qMRI biomarkers and their ability to correctly predict the risk and rate of MCI to AD conversion. Subsequent sections present the data preparation, the utility of the unified approach for the comparison of ML models, and the role of the top qMRI features associated with MCI to AD conversion.

## II. MATERIALS AND METHODS

### A. ADNI/TADPOLE

This study is based on the TADPOLE challenge “standard” data sets (https://tadpole.grand-challenge.org). The TADPOLE sets were derived from the ADNI study (adni.loni.usc.edu). The ADNI was launched in 2003 as a public-private partnership, led by Principal Investigator Michael W. Weiner, MD. The primary objective of ADNI has been to test whether MRI, PET, other biological markers, and clinical and neuropsychological assessment can be combined to measure the progression of mild cognitive impairment (MCI) and early Alzheimer’s disease (AD). For up-to-date information, see www.adni-info.org.

### B. Material

The ADNI/TADPOLE challenge datasets considered for this study were: “D1 – a comprehensive longitudinal data set for training”, and “D2 – a comprehensive longitudinal data set on rollover subjects for forecasting”. The challenge included 1737 individuals from the ADNI database with longitudinal observations. Each subjects’ data included the diagnosis status, neurocognitive evaluations, qMRI longitudinal observations, PET studies, APOE4 status [39]. Detailed information regarding the rational and the information contained in the TADPOLE challenge can be found elsewhere [39]. For this study, we used sex, APOE4, and the 346 longitudinal qMRI measurements provided by the University of California San Francisco (UCSF). UCSF used FreeSurfer Version 4.4, for the analysis of the MRI data sets [40]. The dataset included 864 MCI diagnosed subjects at baseline. 431 of those MCI subjects did not have the structural qMRI data. The 442 MCI subjects with longitudinal qMRI at the baseline were studied in this paper. Among the studied subjects, 187 patients demonstrated MCI to AD conversion and 255 maintained the MCI diagnosis during the observation period. Furthermore, we used normal control patients from the TADPOLE/D1-D2 dataset with qMRI information (n=233) as reference controls. Figure 1 shows the selection process and the main features considered in this paper. Table I shows the demographics of the three groups. The age and gender of the three groups were statistically similar (p < 0.05). As expected, the distribution of APOE4 was statistically different between the MCI to AD converters and those with no conversion (p<0.001).

**Fig. 1.**
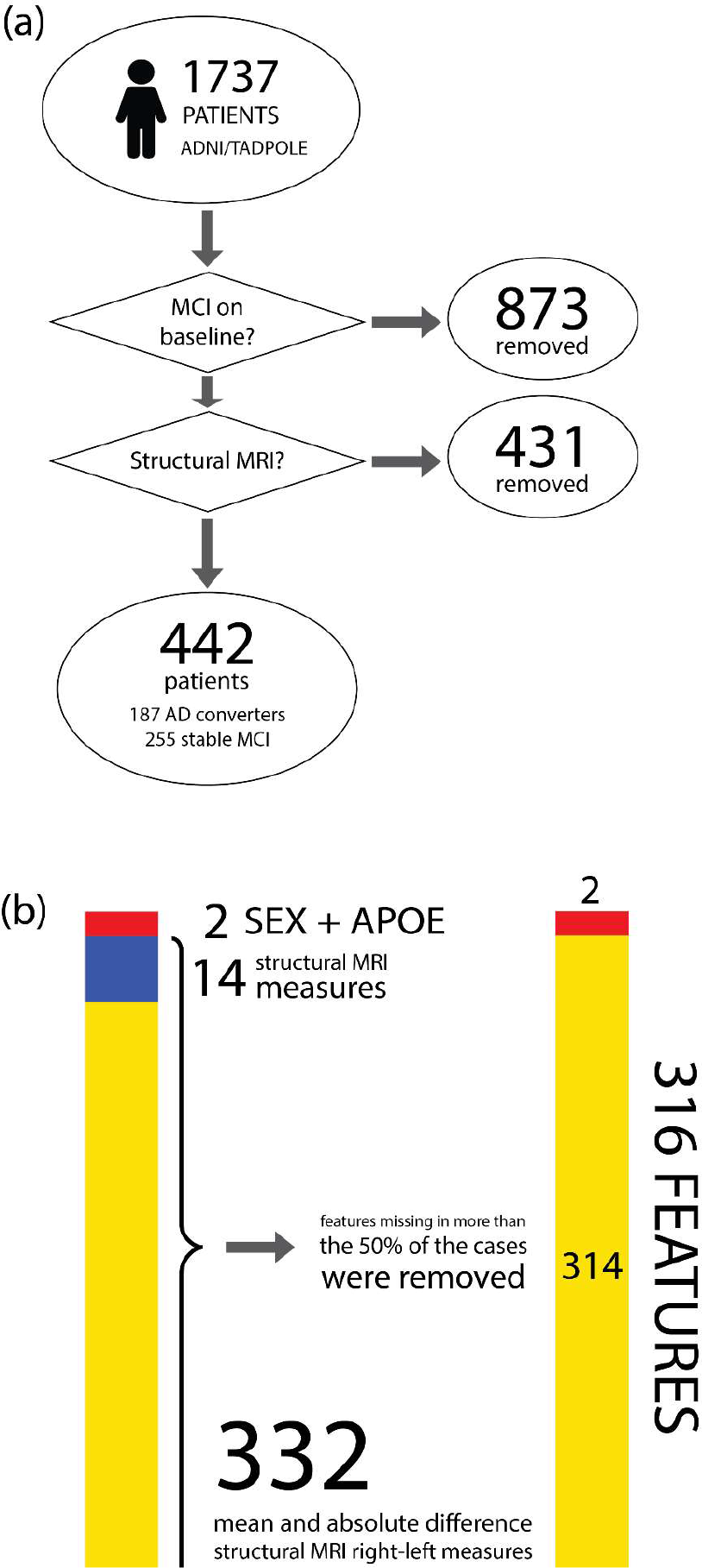
(a) Patient selection process. (b) Feature types used in this study.

**TABLE I.** CHARACTERISTICS OF TADPOLE CHALLENGE SUBJECTS USED IN THIS STUDY. 187 PATIENTS PRESENTED THE MCI TO AD CONVERSION EVENT AND 255 MAINTAINED THE MCI DIAGNOSIS DURING THE OBSERVATION PERIOD. THE NORMAL CONTROL PATIENTS (N=233) WERE USED AS REFERENCE CONTROLS

|  | Sex |  | 1 | Mean<br>Time to<br>event<br>(s.d.) | APOE |  |  |
| --- | --- | --- | --- | --- | --- | --- | --- |
|  | M | F |  |  | 1 | 2 | 3 |
| **MCI to AD** | 107 | 80 | 73.41<br>(7.13) | 848.56<br>(678.96) | 64 | 98 | 25 |
| **No Event** | 150 | 105 | 73.1<br>(7.60) | 1470.22<br>(967.97) | 149 | 81 | 25 |
| **Normal Control** | 113 | 120 | 74.58<br>(5.27) | NA | 167 | 62 | 4 |

### C. Data conditioning and preprocessing

We extended the information provided by the TADPOLE Challenge by computing the time to MCI-to-AD conversion. The event time for stable MCI subjects consisted of the difference in days between the date of the baseline and the date of the last recorded follow-up visit. The event time for subjects that suffer the MCI-to-AD conversion consisted of the difference in days between the date of first AD diagnosis and the baseline date. MCI stable subjects were labeled as censored. After computing the event time, we explored the 346 baseline-qMRI measurements. 332 of these correspond to measures of the left and the right side of the same brain region. Because AD affects both sides of the brain, we described the left-right paired measurements as the mean and absolute differences between them. After that, all the measurements were z-normalized using the 233 normal subjects as reference controls. Finally, qMRI features that were not measured in more than half of the subjects were removed (n=28). After that, the non-reported values of the 314 qMRI features that had majority representation were imputed by the nearest neighbor strategy [41]A complete graphical summary of the data conditioning process can be found in fig 1(b). Fig 2 shows an overall heatmap representation of the analyzed data.

**Fig. 2.**
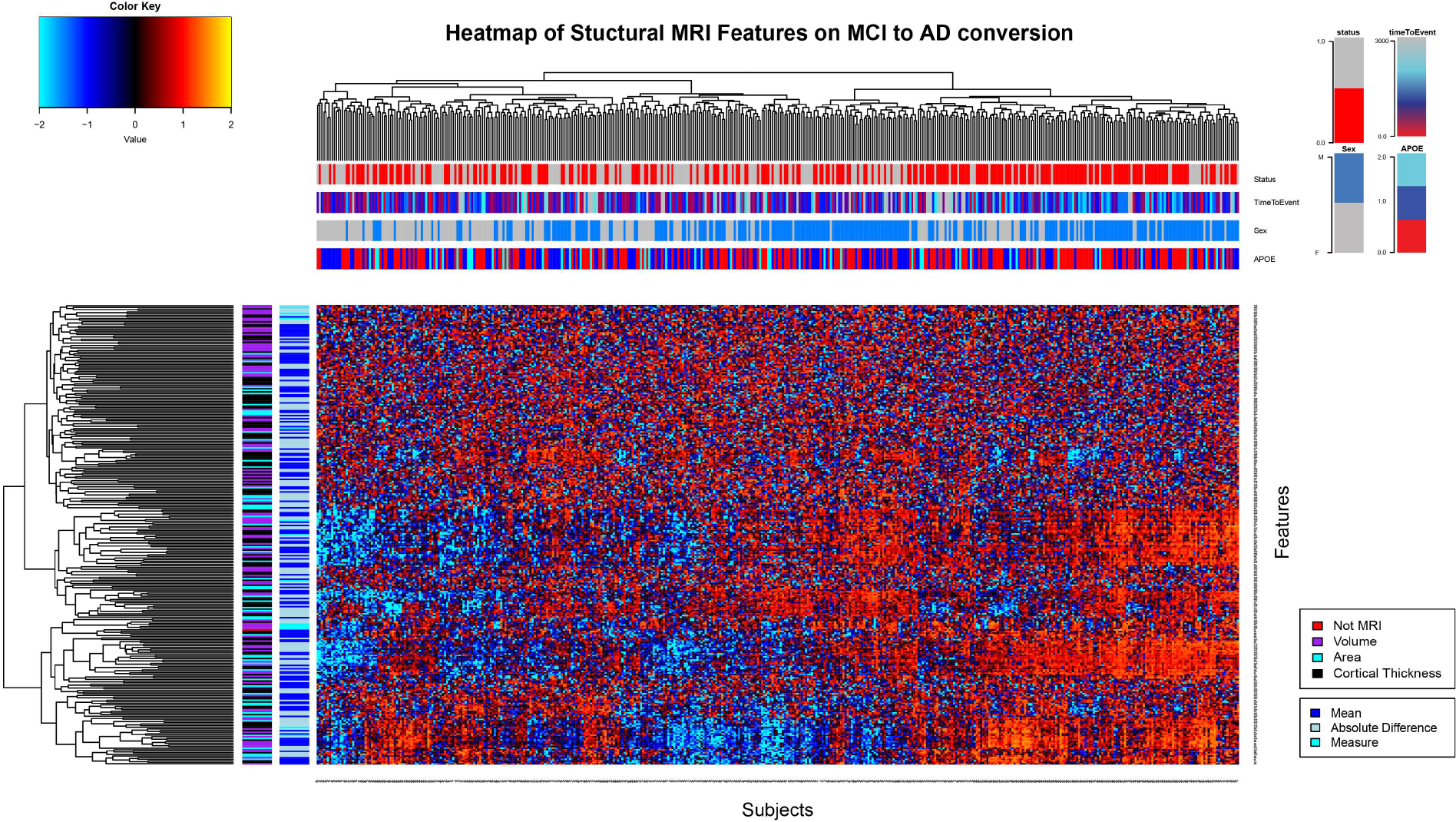
Heat map with 301 features selected by all the Machine Learning Methods. On the top section, patients dendrogram and 4 bars with the subjects’ information about conversion, time to event, sex and APOE. On the left section, dendrogram of features and the information about the type of feature. Subject identification x-axis, features on y-axis.

### D. Machine Learning Methods

The exploration of the set of features and its association with MCI-to-AD conversion was done by learning Cox Regression models. Cox models explore the relationship between the time to the event and the possible explanatory variables. The model estimates the hazard *λ*_*i*_ of the subject *i* given the observed feature vector ***X*_*i*_** = {*X*_*i***1**_, …, *X*_*ip*_}, and the unknown baseline hazard *λ*(*t*_0_). i.e,

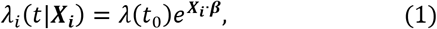

where ***β*** = {*β*_1_, …, *β*_*P*_} is the vector of coefficients. Hence, the Cox model provided an estimate of total hazard (risk) of conversion, for an individual, given the observed features. Due to the large set of possible qMRI features to be considered in the Cox model, machine learning methods were used to find the “optimal” set of features and their corresponding coefficients that mimiced the observed rate of conversion.

There are several strategies aimed to build Cox models. In this paper, we evaluated three publically available ML methods that build Cox models from a set of features, and we compared the results to a simple Cox-based univariate filter. The first method was the Bootstrapped Stage-wise Model Selection (BSWiMS). BSWiMS is part of the FRESA.CAD R package and is a supervised model-selection method aimed to select a unique statistical model that predicts a user-specified outcome, in this case, a survival outcome. The statistical model is constructed by bagging a set of Cox models built by the unique set of model-wise statistically-significant features [36]. The second method was the Penalized Cox Regression (CoxNet) and part of the gmlnet R package. This algorithm fits the Cox Model regularized by an elastic net penalty [34]. It was executed in LASSO mode with cross-validation to determine the value of lambda that returned the smallest error. Briefly, the LASSO mode considers the L1 regularization only, which decreases the coefficients by a constant (lambda) to perform feature selection removing those coefficients lower than lambda. The third method used was the Primal-dual Active set. This method is part of the BeSS (Best subset selection) R package. This method uses an efficient active set algorithm to choose the best possible Cox model. We executed this method with the sequential search strategy [35]. Furthermore, we used the features returned by the models-selection methods as a filter strategy for fitting Cox models. Finally, in the fourth strategy, we explored the univariate Cox analysis to filter-out no statistical significant features inside the Cox models from the above methods. The p-value of the univariate fit was adjusted for false discovery rate (FDR) [42]. The coefficients whose adjusted p-value were smaller than a user-supplied threshold were included in the final CoxPH model. All filter-based Cox-models were fitted using CoxPH (CoxPH Model from Survival R Package). In summary, we evaluated six different methods for building Cox-Models: LASSO, BSWiMS, BeSS, and three more filtering features from all these.

#### 1) ML Evaluation and Validation

The main aim of this paper is the comprehensive evaluation of different ML approaches that return or select “optimal” Cox models. We used repeated holdout cross-validation (RHOCV), for the evaluation of different ML strategies. The test results of the RHOCV were used to compare and explore the performance of the machine learning alternatives. The RHOCV strategy was implemented as an extension of FRESA.CAD R package^2^

The strategy divided the data into random training and testing sets with a user-supplied train fraction. The training set was used for model selection, while the holdout set was used to validate the trained method [36]. Furthermore, the RHOCV implementation used the R package *Survival* to calculate the final Cox predictions of each selected model. Cox predictions returned the linear predictions, the risk, and the expected follow-up times.

Our implementation of RHOCV returned the execution times, the Jaccard index, the model size, and the training and testing samples of every single method. The Jaccard index computed the average similarity between the selected features between models, and can be written as:

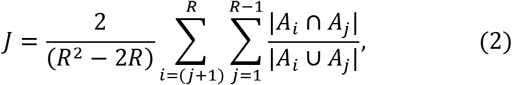

where *R* is the number of holdout repeats, and *A*_*j*_ is the set of the *k* selected features for the Cox model of the *j* holdout training sample. The range of the index varies from 0 to 1, where 1 represents that the feature selection method always selects the same set of features on each repetition.

The R implementation also reported summary statistics of the test results. The Cox-fitted coefficients ***β***^*j*^ on each training set ***T***_j_ were used to get the linear predictions 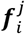 of the holdout set 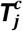 at each repetition:

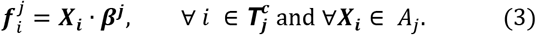

Once all the test predictions were obtained for each repetition, the testing results were summarized by computing the median prediction of each subject: 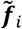 = median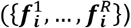. The median prediction was used to divide the groups into: High-risk 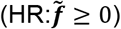 vs Low-risk 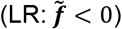. The receiver operating characteristic (ROC) plots and their area under the curve (AUC) with their corresponding 95% confidence intervals (95%CI) were computed for the median prediction using the pROC package [43]. Accuracy (*ACC*), sensitivity (*SEN*), and specificity(*SPE*) describing the ability of the Cox models to predict censored vs uncensored subjects were computed based on the number of true positives (*TP*), and true negatives (*TN*)

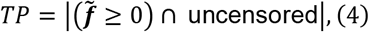

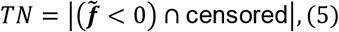

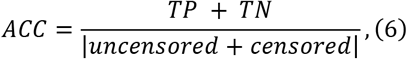

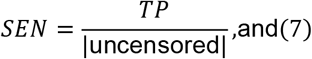

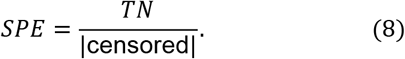

The implemented code also reported the 95%CI of *SEN*, *SPE*, and the *ACC*. Furthermore, the concordance index (c-index) of expected times vs to follow-up times, also was analyzed. The c-index is a performance measure of survival models and is the fraction of all order pairs of subjects *ε*_*ij*_ whose predicted survival times are correctly ordered among all subjects that can actually be ordered. It can be written as:

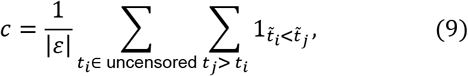

where the indicator function 1_a<b_ = 1 if *α* < *b*, and 0 otherwise. |*ε*| is the number of ordered pairs. 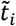 is the median of the predicted survival time, and *t*_*i*_ is the actual observed time of the uncensored subject *i*. The values of the concordance index range from 0 to 1, where 1 implies a perfect concordance between observed and predicted times.

The visualization of the predicted survival (HR vs LR) groups was done using Kaplan-Meier plots of survminer R package [44]. The statistical significance of the difference between the two survival groups was evaluated by the Logrank test [45].

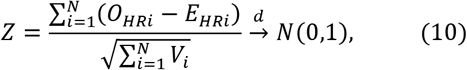

where *i* is the rank of the event-time ordered population, *O*_*HRi*_ represent the actual observed events, *E*_*HRi*_ = (*O*_*i*_/*N*_*i*_)*N*_*HRi*_ the expected number of events, and *V*_*i*_ is the total variance,

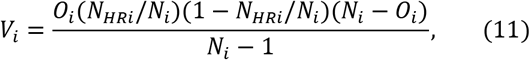

where *O*_*i*_ is the actual of number of events, *N*_*i*_ is the number of subjects below rank, and *N*_*HRi*_ is the number of subjects at HR. We ran the RHOCV 20 times. Each run used 70% of the subjects (n=309) for training the other 30% (n=133) for testing. These settings produced, on average, 6 test predictions per subject.

## III. RESULTS

Table II shows the main results of the RHOCV on the six tested models. We report the major findings per method and all the performance statistics with 95% CI.

**TABLE II.**
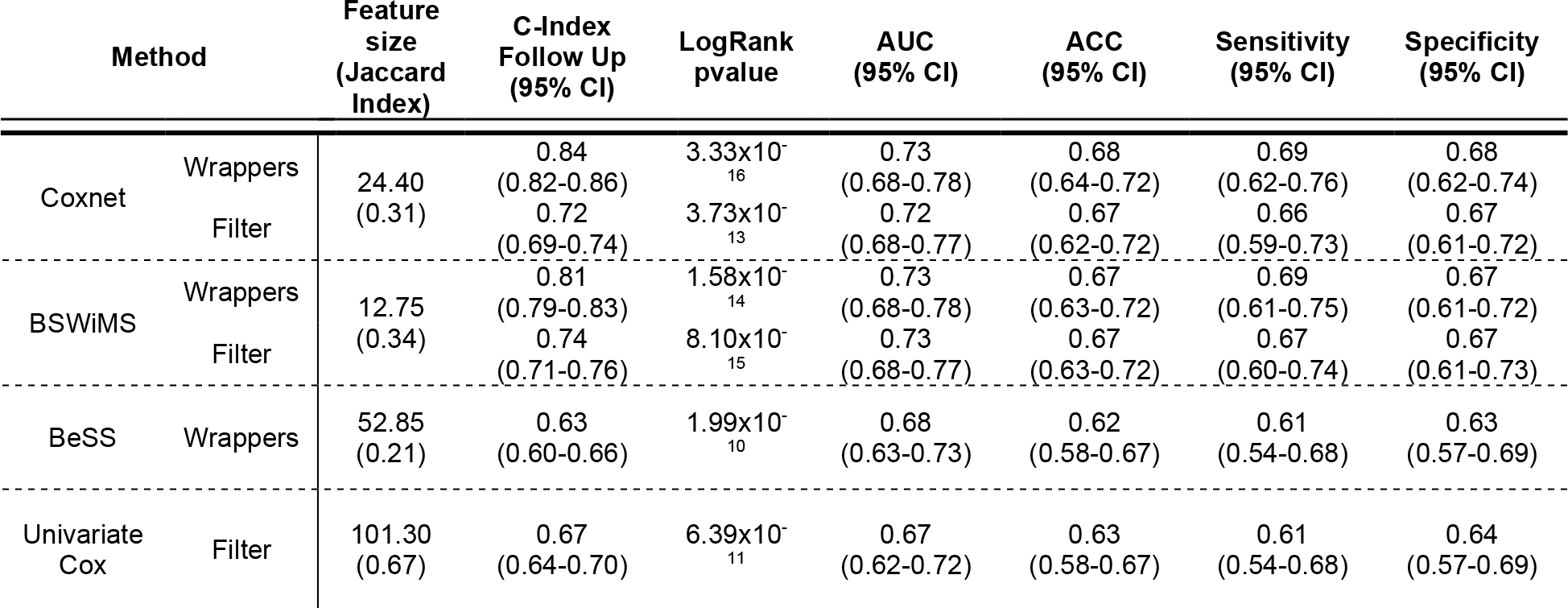
MODELS PREDICTIONS STATISTICS. CONTAINS C-INDEX OF FOLLOW-UP TIMES PREDICTIONS, THE P-VALUE ON LOG RANK TEST BETWEEN LOW-HIGH RISK CURVES, AREA UNDER THE CURVE, ACCURACY, SENSITIVITY AND SPECIFICITY WITH THEIR 95% CONFIDENCE INTERVALS

The BSWiMS strategy selected the smallest models. They contained an average of 13 features with an average Jaccard index of 0.34. The mean volume of the amygdala and entorhinal and the mean cortical thickness average of bankssts were selected on every iteration. The BSWiMS model had c-index of 0.81 (0.79-0.83) with *ACC* = 0.67 (0.63, 0.71), *SEN* = 0.69 (0.61,0.75), *SPE* = 0.67 (0.61,0.72), and *AUC* = 0.73 (0.68,0.78). The Cox Modeling based on BSWiMS reported the following classification performance: *ACC* = 0.67 (0.63, 0.72), *SEN* = 0.67 (0.60, 0.74), *SPE* = 0.67 (0.61, 0.73), and *AUC* = 0.73 (0.68, 0.77). Hence BSWiMS models were very similar to CoxPH fitted model. Figure 3(a) shows the Kapplan-Meier curves of the subjects predicted at risk of conversion vs the subjects predicted as stable for the Cox model created by BSWiMS features.

**Fig. 3.**
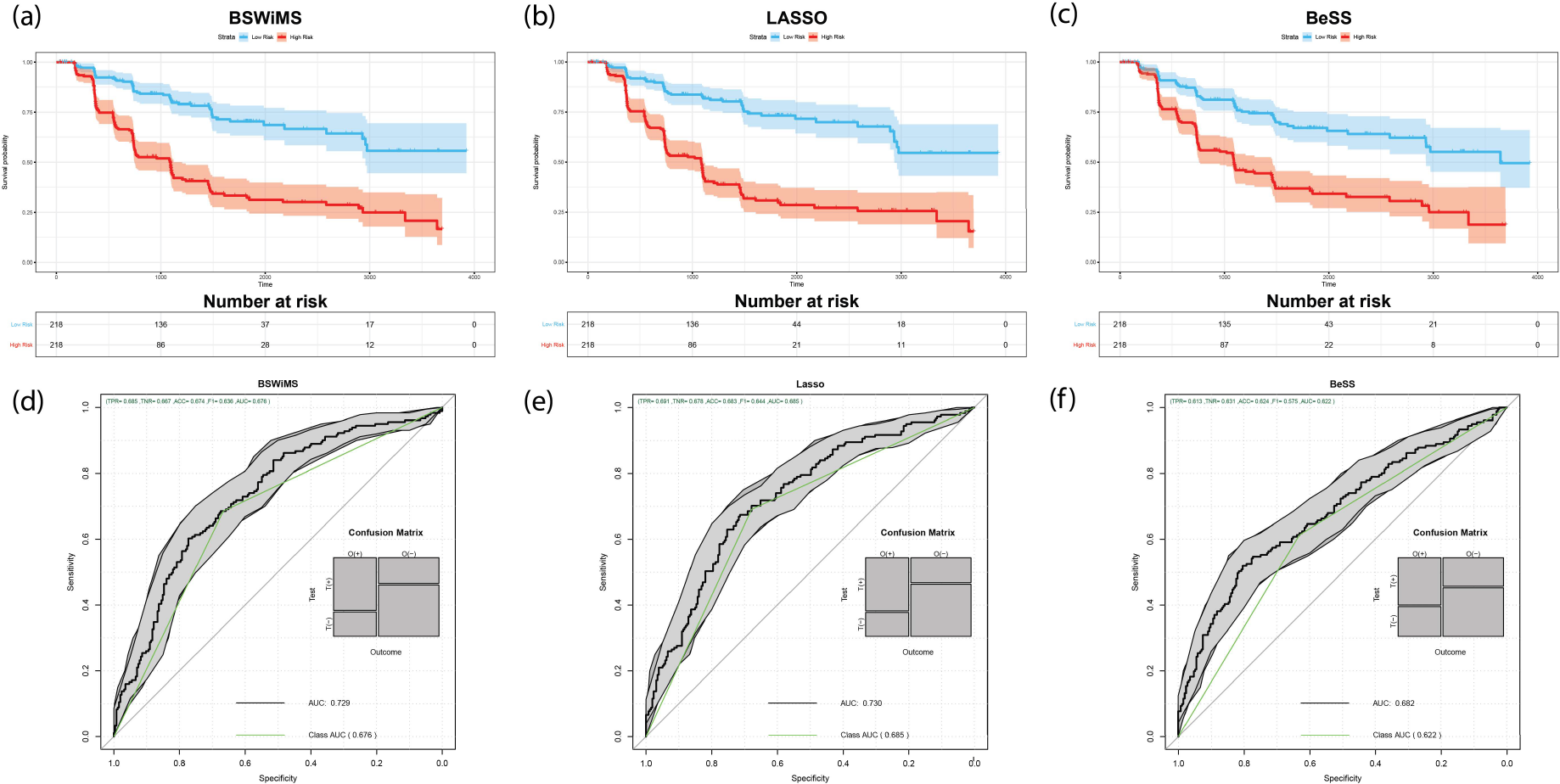
Kaplan Meier (KM) and ROC curves for wrappers/embedded section. CoxNet showed the best accuracy on the classification and the best c-index on Risk and Follow-up times. (a) Model 1 BSWiMS KM (b) CoxNet KM (c) BeSS KM (d) BSWiMS ROC (e) CoxNet ROC (f) BeSS ROC

The CoxNet/LASSO method generated models with an average set of 24 features with a Jaccard index of 0.28. The most common features were APOE4, the mean cortical thickness average of Bankssts and the mean volume (cortical parcellation CP) of entorhinal. 75% of the repetitions selected the mean volume (cortical parcellation) of inferior temporal, the absolute difference of cortical thickness average of pars opercularis, the mean volume (WM parcellation) of the amygdala and the mean cortical thickness standard deviation of bankssts. This model reported c-index = 0.84 (0.82-0.86), *ACC* = 0.68 (0.64, 0.73), *SEN* = 0.69 (0.62, 0.76), *SPE* = 0.68 (0.62, 0.74), and *AUC* = 0.73 (0.68, 0.78). The Cox regression models fitted with LASSO features returned the following performance: *ACC*= 0.67 (0.62, 0.71), *SEN* = 0.66(0.59, 0.73), *SPE* = 0.67 (0.60, 0.72) and *AUC* = 0.72 (0.68, 0.77). These results indicate that CoxPh performance is lower than L1 fitted model, implying that L1 penalization helped in improving the prediction of which subjects converted. Figure 3(b) shows the Kapplan-Meier curves.

The BeSS method returned on average models with 53 features with a Jaccard index of 0.21. Three features were selected on every single repetition: APOE4, mean cortical thickness standard deviation of bankssts and mean volume (cortical parcellation) of entorhinal. 75% of the time the following 3 features were selected: mean cortical thickness standard deviation of temporal pole, mean cortical thickness standard deviation of the rostral middle frontal and mean surface area of cuneus. BeSS models reported c-index = 0.63 (0.60, 0.66), *ACC* = 0.63 (0.58, 0.67), *SEN* = 0.61 (0.54, 0.68), *SPE* = 0.63 (0.57, 0.69), and *AUC* = 0.68 (0.63,0.73).

Finally, the models created by univariate Cox filter were the largest. The average size of the models included 103 elements with a Jaccard index of 0.65. 54 features were selected in all the iterations. Among the selected features were APOE4, the mean cortical thickness average of Parahippocampal, the cortical thickness average and the volume (cortical parcellation) of pars opercularis. Classification performance of univariate filter were: *ACC* = 0.63 (0.58, 0.67), *SEN* = 0.61 (0.54,0.68), *SPE* = 0.64 (0.57,0.69) and *AUC* = 0.67 (0.62,0.72). Hence the Cox models based on simple univariate filter had the least robust performance. Fig 3 and Fig 4 show the complete Kapplan-Meier curves and the ROC plots based on the median estimations for ML-methods and filter-based-methods.

**Fig. 4.**
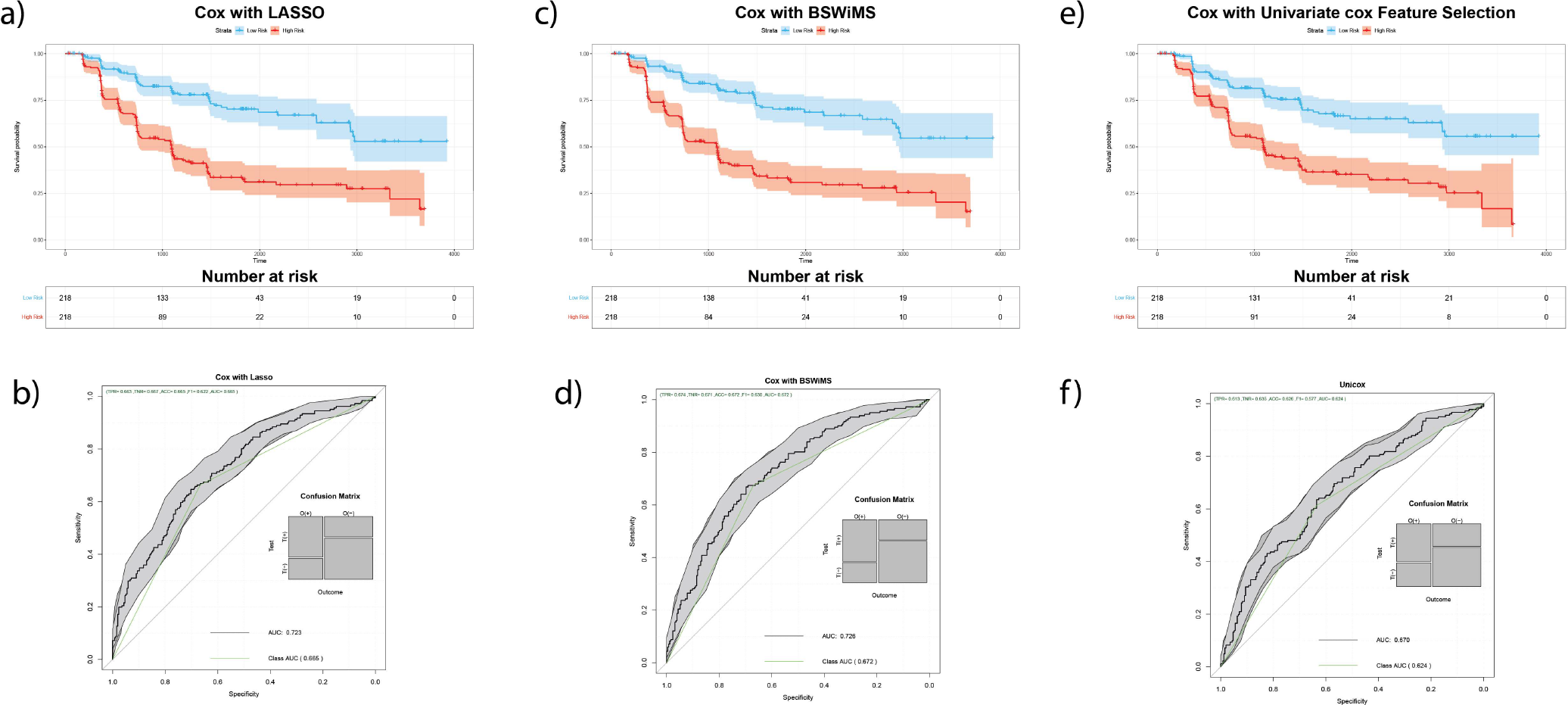
Kaplan Meier (KM) and ROC curves for filters section. Cox Model build with BSWiMS features showed the best accuracy on the classification and the best c-index on Risk and Follow-up times. (a) Model 4 Cox with BSWiMS KM (b) Cox with BSWiMS ROC (c) Model 5 Cox with CoxNet KM (d) Cox with CoxNet ROC (e) Model 6 Cox with Univariate Cox KM (f) Cox with Univariate Cox ROC

We performed a detailed analysis of the set of selected features across ML methods. The analysis of the RHOCV reported that ten features were common on 50% of the sets. To evaluate the importance of these ten features as a risk factor for MCI to AD conversion, we refit the Cox model using these ten features. We then reported the hazard ratios (HR) and their corresponding 95% CI: The mean volume (CP) of entorhinal HR = 0.63 (0.50, 0.80), mean cortical thickness SD of Bankssts HR = 1.60 (1.20,2.12), APOE4 HR = 1.74 (1.40,2.17), mean volume (WMP) of amygdala HR = 0.88 (0.69,1.13), mean cortical thickness AVG of Bankssts HR = 0.76 (0.60,0.97), mean volume (CP) of inferior temporal HR = 0.79 (0.62,1.00), absolute difference cortical thickness AVG of middle temporal HR = 1.38 (1.07, 1.78), absolute difference of cortical thickness AVG of pars opercularis HR = 1.46(1.09, 1.94), absolute difference cortical thickness average of inferior parietal HR = 1.33 (1.04, 1.70), mean cortical thickness standard deviation of Rostral middle frontal HR = 0.64(0.50, 0.81). A heatmap representation with the ten features correlation with the outcome can be found in Figure 5. Table III provides more details of the ten characteristics. The last two columns of table III shows the rank of the features of the four ML approaches. The MV HR and the UV HR correspond to the Hazard ratios of the feature inside a Multivariate model and the HR computed by the univariate approach respectively. It is clear that feature ranking and importance depended on the ML method.

**Fig. 5.**
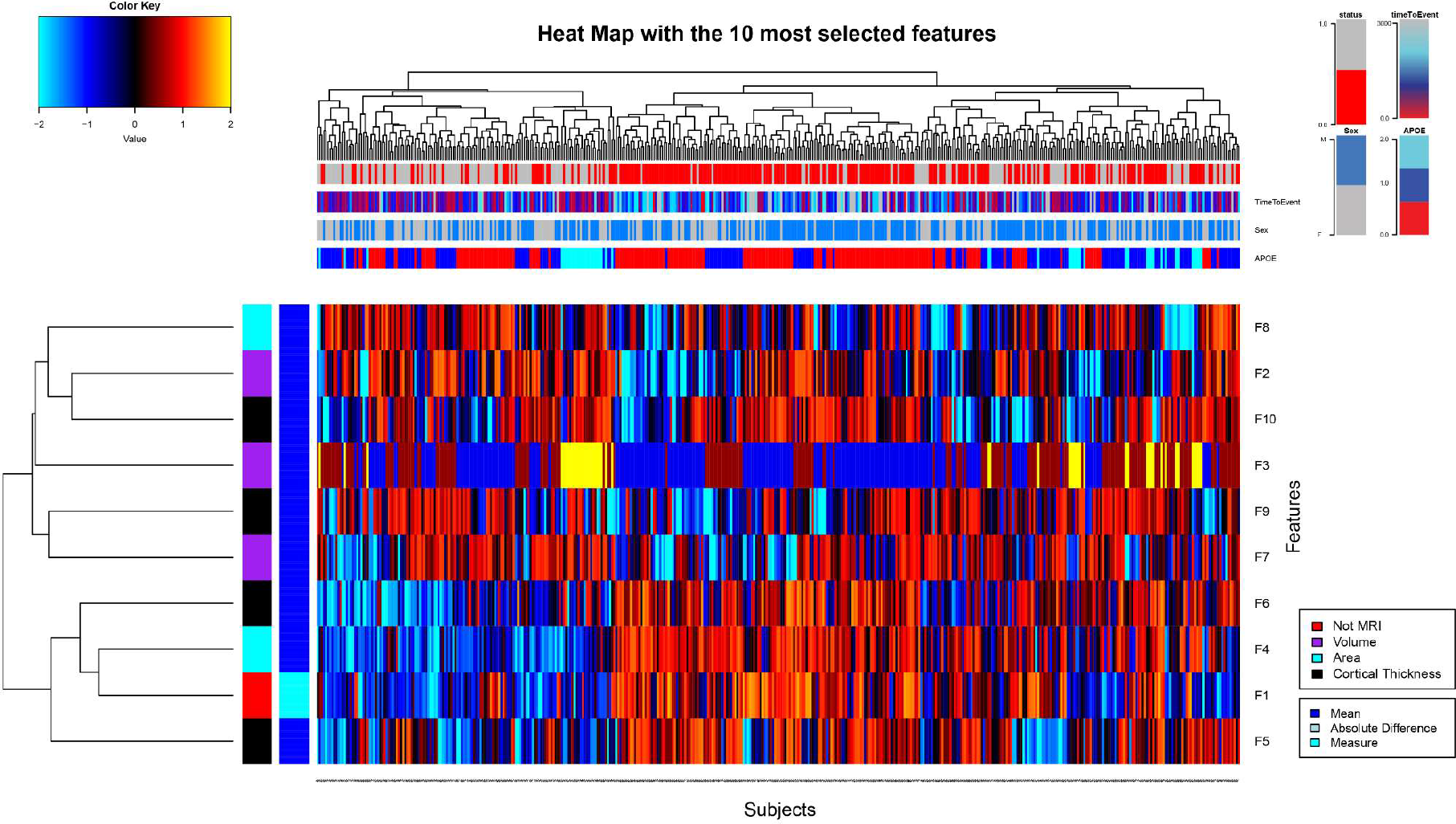
A heat map representation of the features associated with MCI to AD conversion. The figure shows the ten features selected by all the 4 methods in at least in the half of the iterations (horizontal axis) and subjects on the vertical axis. (F1) Mean volume (CP) of entorhinal, (F2) mean cortical thickness SD of Bankssts, (F3) APOE4, (F4) mean volume (WMP) of amygdala, (F5) mean cortical thickness AVG of Bankssts, (F6) mean volume (CP) of inferior temporal, (F7) absolute difference cortical thickness AVG of middle temporal, (F8) absolute difference of cortical thickness AVG of pars opercularis, (F9) absolute difference cortical thickness AVG of inferior parietal, (F10) mean cortical thickness SD of Rostral middle frontal

**TABLE III.**
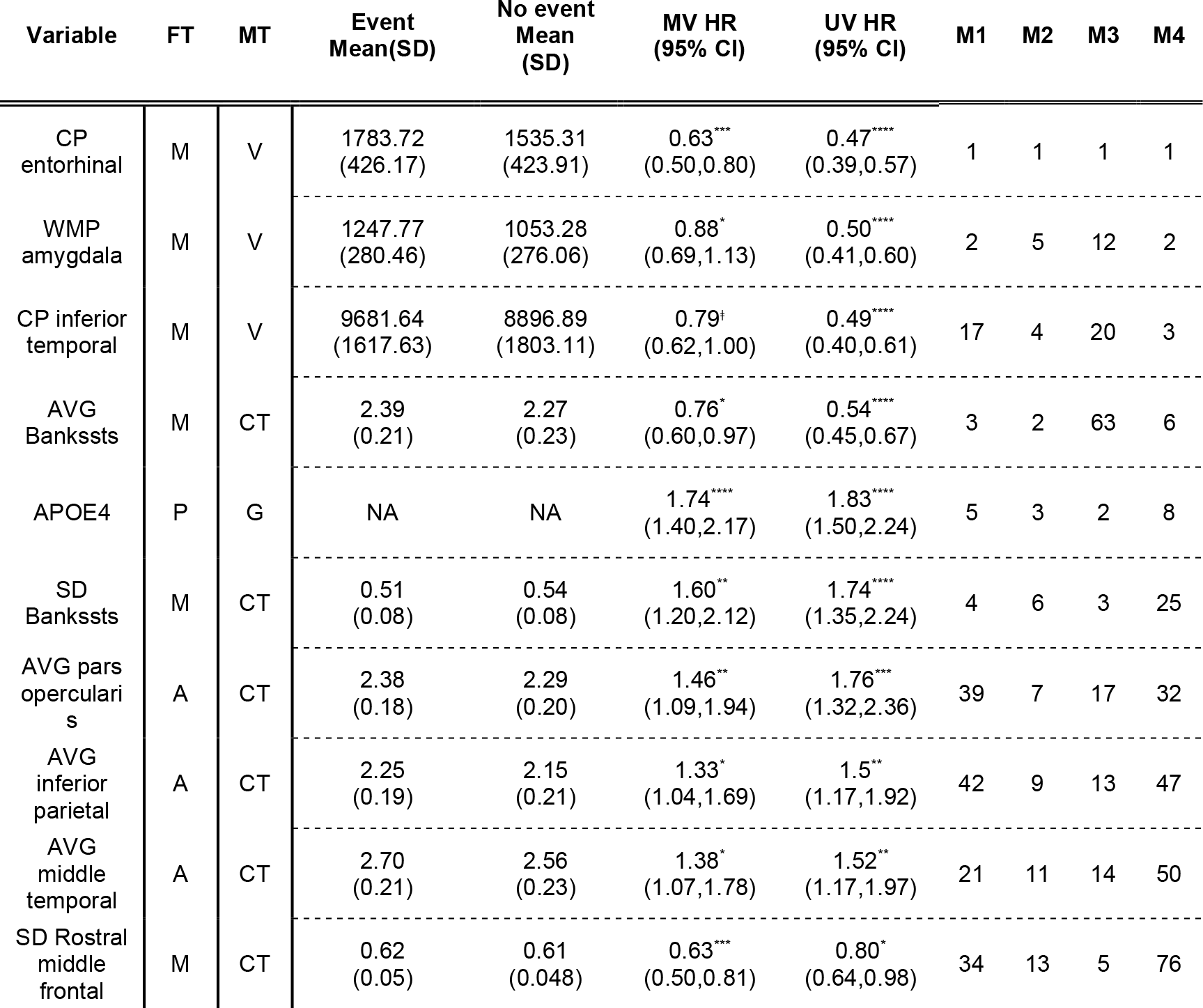
CHARACTERISTICS AND RANKING OF TEN FEATURES SELECTED IN ALMOST THE HALF OF THE ITERATIONS. THE RANKING WAS ORDERED BASED ON THE NUMBER OF TIMES SELECTED AND THEN ORDERED DEPENDING ON THE P-VALUE OF UNIVARIATE COX ANALYSIS. [FT = FEATURE TYPE; M=MEAN, P= POLYMORPHISM, A=ABSOLUTE DIFFERENCE], [MT = MEASURE TYPE; V=VOLUME (MM3), G = GENE, CT = CORTICAL THICKNESS (MM)], [M1 = BSWIMS, M2 = COXNET/LASSO, M3 = BESS, M4 = UNIVARIATE COX] P. VALUE SIGNIFICANCE: ǂ <0.1, * <0.05, ** <0.01, *** <0.001, **** <10-04

## IV. DISCUSSION

In this work, we compared four different ML strategies that generated six proportional hazard models from qMRI structural analysis of MCI patients that either converted to AD or remained as MCI. The first three strategies – BSWiMS, LASSO, and BeSS – returned a Cox regression model and the set of features that were required to make an accurate estimation of the risk of conversion. The fourth strategy was a filter approach; hence selected features were used to build a standard Cox regression model. This last strategy was evaluated with the features generated by the first three methods and the features generated by a univariate Cox regression model. The performance of six proportional hazard models was evaluated using RHOCV and the most common features analyzed to report their importance in the rate of MCI to AD conversion.

### A. ML Method Validation

The RHOCV evaluation created 20 random splits of the dataset into training and test set. For each such split, the train fraction was 0.7 and the remaining 0.3 was used for testing.This evaluation strategy allowed the evaluation of the effect of the training set on feature selection, and, at the same time, permited a training-set unbiased evaluation of the test performance. The reported results indicated that ML methods selected models with very different internal features. Model sizes varied from method to method and ranged from a minimum of 13 features to complex multivariate modeling based on 103 features. The six qMRI-based models reported c-index ranging from 0.63 to 0.84. The simplest model overperformed the most complex one: 0.84(CI 0.82,0.86) for Coxnet vs 0.63(CI 0.60,0.66) for BeSS.

Regarding this classification performance, it is important to note that proportional hazard models were not designed for classifications task. To address this issue, we assumed that subjects predicted to have an increased risk of conversion (Risk > 1) should correspond to true MCI to AD conversion, while subjects at low-risk prediction (Risk <= 1) should correspond to MCI-stable subjects. This strategy allowed us to evaluate the accuracy, sensitivity, specificity, and *AUC* of the Cox regression models. The reported *AUC* performance of the methods ranged from 0.67 to 0.73 for their potential to detect patients at risk of conversion. This performance was slightly lower to other methods based on SVM or Logistic Regression classifiers [46]. To test the impact of using all subjects in ROC AUC analysis, we conducted a post hoc experiment. In this experiment, we analyzed test prediction on MCI stable subjects whose last visit was greater than 4 years (146 no-event subjects did not meet the criteria). This change in selection criteria resulted in the ROC curve presented by Figure 6. We clearly see that Cox based conversion risk prediction had a similar performance (ROC ACU= 0.79) to previous works [47].

**Fig. 6.**
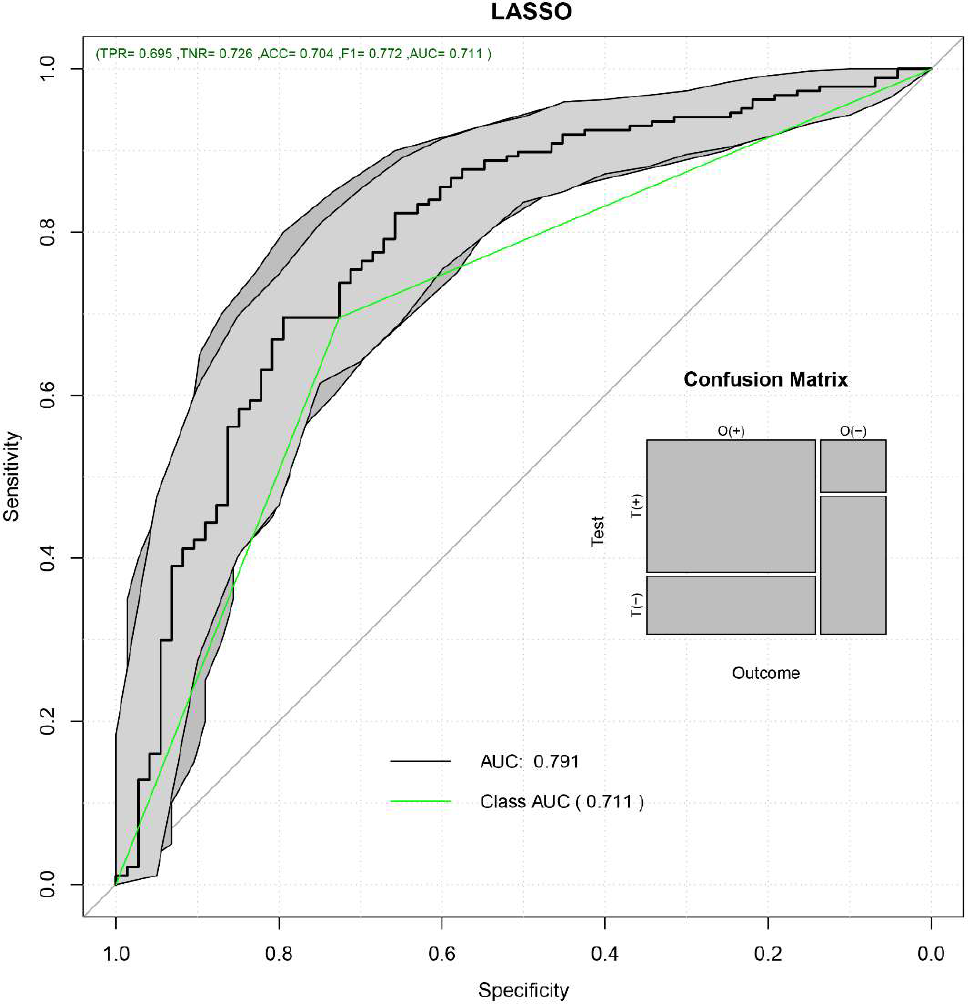
Coxnet ROC with 296 patients who suffered the conversion or have a censored event in more than 4 years.

Regarding the reproducibility of feature selection, the Jaccard analysis indicated that the internal structure of the Cox models depended on the training set. The method with the largest Jaccard index (0.65) was based on the univariate filter, and also was the method with the largest set of features and with the poorest performance. The smallest models were returned by the BSWiMS strategy. It had a Jaccard Index 0.35, implying that only 35% of the features overlapped across different training sets. These results put forward that the discovery of risk factors associated with MCI to AD conversion depended on the training set and the machine learning strategy used to discover risk factors. This observation is supported by the literature, where different authors have reported a different set of features associated with MCI to AD conversion.

### B. Feature Relevance and Analysis

This work analyzed 316 features and their role in MCI to AD conversion risk. The RHOCV reported that 301 out of the 316 characteristics may have some association, but the detailed analysis indicated that only ten features were selected at least 50% of the time. Many of these ten features have already been reported as potential biomarkers associated with MCI to AD conversion [8], [28], [29], [48]. APOE4, a factor that has been validated several times as a biomarker indicative of the risk of conversion [8] was an important validation in our work. qMRI related features included the decreased volume of the cortical parcellation of the entorhinal, the increase in the white matter parcellation of the amygdala and increase volume and thickness standard deviation of Bankssts. These findings confirmed the results of previous studies [28], [48] [49]. Regarding novel features, our work suggests that large differences between left-right brain structures like the Pars Opercularis, Middle Temporal Lobule, and the Inferior Parietal Lobule, unlike the volumes listed for each condition and structure as mentioned in previous studies [50]. [51], [52].

This work also aimed to improve the knowledge of the role of these features in the AD process; hence we reported the list of the top biomarkers along with their standardized HR associated with the conversion of MCI to AD. Reporting HR per z-units of the normal distribution may help physicians predict how far a specific patient in their MCI to AD process is.

### C. Limitations

The results presented in this work are limited in three key aspects. First, patient misdiagnosis is present, hence affecting feature selection and model building. The dementia diagnosis of “true” AD patients is not an exact science, hence detecting the exact time of conversion is also prone to diagnoses errors, and these two errors are present in modeled survival outcome. Second, it is based on the ADNI cohort and measurements; therefore, it is biased towards the environmental factors present in the US and the Caucasian race. Third, we assumed that all MCI will convert to AD in some point in the future. This assumption should not be a major issue if the proportion of misdiagnosed MCI is low. These key limitations indicate that the presented findings have to be confirmed on cohorts from different countries and ethnicities.

## V. CONCLUSION

ML is a viable strategy to build and explore survival models from hundreds of candidate features. This work presented a comprehensive evaluation of four ML strategies for building Cox regression models that predicted the time from MCI to AD conversion based on the qMRI analysis provided by ADNI. The evaluation included associations to event time, risk classification performance, and detailed characterization of the main features associated with MCI to AD conversion. The reported findings indicate that Cox-based discovery depends on the subjects selected for training as well as the machine learning method used for model selection.

## ACKNOWLEDGMENT

This work was partially supported by Secretaría de Educación Superior, Ciencia, Tecnología e Innovación part of Gobierno de la República del Ecuador and by Strategic Research Group of Bioinformatics for Clinical Diagnosis from Tecnólogico de Monterrey. Data collection and sharing for this project was funded by the Alzheimer’s Disease Neuroimaging Initiative (ADNI) (National Institutes of Health Grant U01 AG024904) and DOD ADNI (Department of Defense award number W81XWH-12-2-0012). ADNI is funded by the National Institute on Aging, the National Institute of Biomedical Imaging and Bioengineering, and through generous contributions from the following: AbbVie, Alzheimer’s Association; Alzheimer’s Drug Discovery Foundation; Araclon Biotech; BioClinica, Inc.; Biogen; Bristol-Myers Squibb Company; CereSpir, Inc.; Cogstate; Eisai Inc.; Elan Pharmaceuticals, Inc.; Eli Lilly and Company; EuroImmun; F. Hoffmann-La Roche Ltd and its affiliated company Genentech, Inc.; Fujirebio; GE Healthcare; IXICO Ltd.; Janssen Alzheimer Immunotherapy Research & Development, LLC.; Johnson & Johnson Pharmaceutical Research & Development LLC.; Lumosity; Lundbeck; Merck & Co., Inc.; Meso Scale Diagnostics, LLC.; NeuroRx Research; Neurotrack Technologies; Novartis Pharmaceuticals Corporation; Pfizer Inc.; Piramal Imaging; Servier; Takeda Pharmaceutical Company; and Transition Therapeutics. The Canadian Institutes of Health Research is providing funds to support ADNI clinical sites in Canada. Private sector contributions are facilitated by the Foundation for the National Institutes of Health (www.fnih.org). The grantee organization is the Northern California Institute for Research and Education, and the study is coordinated by the Alzheimer’s Therapeutic Research Institute at the University of Southern California. ADNI data are disseminated by the Laboratory for Neuro Imaging at the University of Southern California.

## Footnotes

2 https://github.com/joseTamezPena/FRESA.CAD

